# Interleaved Replay of Novel and Familiar Memory Traces During Slow-Wave Sleep Prevents Catastrophic Forgetting

**DOI:** 10.1101/2025.06.25.661579

**Authors:** Ryan Golden, Rajat Saxena, Oscar C. González, J. Erik Delanois, Scott Kilianski, Bruce L. McNaughton, Maxim Bazhenov

## Abstract

Humans and animals can learn continuously, acquiring new knowledge while integrating it into a lifelong memory pool. In contrast, artificial neural networks (ANNs) suffer from catastrophic forgetting, where new training disrupts existing memories. This issue can be alleviated in ANNs by interleaving training on new tasks with past data; however, whether the brain uses a similar strategy is unknown. In this work, we show that slow-wave sleep interleaves replay of familiar and novel (i.e. hippocampal-dependent) memory traces within individual slow waves, allowing new memories to integrate into the existing cortical pool without interference. This study presents a novel theory for how memory traces acquired across an animal’s life are organized within the cortical-hippocampal system to support continual learning and suggests novel principles for a broad range of continual learning AI.

## INTRODUCTION

Human and non-human animals alike have a remarkable ability to learn continuously and from relatively few examples, incorporate new data into their corpus of existing knowledge, and generalize episodic memories beyond a single experience. In contrast, artificial neural networks (ANNs) suffer from “catastrophic forgetting” whereby they achieve optimal performance on newer tasks at the expense of performance on previously learned tasks ^1–4^. The dichotomy between learning new tasks and the ability to retain and generalize knowledge across all tasks is characterized by the stability–plasticity trade-off ^2,5,6^. On the one hand, a network must be plastic such that the parameters in the network can change in order to accurately represent and respond to new tasks. On the other hand, a network must be stable such that it maintains knowledge of older tasks.

A key proposal in the connectionist literature was the Complementary Learning Systems (CLS) framework ^1,7^. CLS posits two complementary systems: a fast-learning hippocampal system that stores each new episode and a slow-learning cortical system that integrates information gradually. Offline replay - especially during sleep - allows the fast system to repeatedly present novel memory traces to the slow system, effectively interleaving old and new experiences to update the cortical memory system with minimal interference. A critical element of the consolidation stage is cortical replay triggered by hippocampal sharp wave–ripple (SWR) events ^8^. While powerful, CLS alone does not explain how new memories encoded by hippocampal SWRs avoid overwriting cortical traces of overlapping, familiar memories.

Within artificial networks, a proposed solution to catastrophic forgetting is rehearsal ^1,9,10^ or pseudorehearsal ^11^ - that is, interleaving new memory inputs with samples from previous tasks. Since the brain does not store or generate old input samples, it is unlikely to utilize rehearsal directly. However, the brain retains synaptic weights, and the memory traces they encode can be reactivated through synaptic replay. Could sleep provide the biological mechanism that enables interleaved replay of old and new memories without requiring explicit access to old memory samples?

Using a combination of biophysical modeling and analyses of single-unit activity from the mouse retrosplenial cortex – while navigating either a highly familiar environment or a completely novel one - we found that slow-wave sleep interleaves replay of familiar and novel (i.e. hippocampal-dependent) memory traces within individual slow waves. Hippocampal SWRs arriving near the Down-to-Up or Up-to-Down transitions of the sleep slow oscillation (SO) can entrain novel memory replay, while the middle phase of the Up state tends to replay familiar cortical memories. This strategy supports the formation of new cortical memory traces while minimizing interference and damage to existing ones and likely depends on bidirectional cortico-hippocampal dialogue ^12–15^. Our study introduces idea of Structured Cortical Replay (SCoRe) - a framework explaining how memory traces acquired at different times during an animal’s lifespan are dynamically organized within the thalamo-cortico-hippocampal system to enable continual learning and offering insights into potential strategies for mitigating catastrophic forgetting in artificial networks.

## RESULTS

### Modeling experiments

The network model utilized throughout the study was built upon thalamocortical models previously used in earlier work ^16,17^. In short (see details in **Methods**), the basic thalamocortical circuit (Figure 1A) consists of a single cortical layer with excitatory pyramidal cells (PYs) and inhibitory interneurons (INs), and a single thalamic layer with excitatory thalamocortical cells (TCs) and inhibitory reticular interneurons (REs). All neurons were modeled using the Hodgkin-Huxley formalism. Synaptic connections were set deterministically within a local radius and held constant, except for PY-PY synapses, which were subject to plasticity. These synapses were set probabilistically within a local radius, with initial weights drawn from a Gaussian distribution (Figure 1B), and were allowed to evolve according to local spike-timing-dependent plasticity (STDP) rules. Transitions between awake and slow-wave sleep (SWS) states were simulated by changing cellular and synaptic parameters to mimic the effects of the distinct neuromodulatory tone of each brain state ^16^. Cortical PYs spiked asynchronously during the awake state, while during sleep their activity was organized by transitions between active (Up) and silent (Down) states of the slow oscillation (SO ^18^; <1Hz) (Figure 1C).

**Figure 1.**
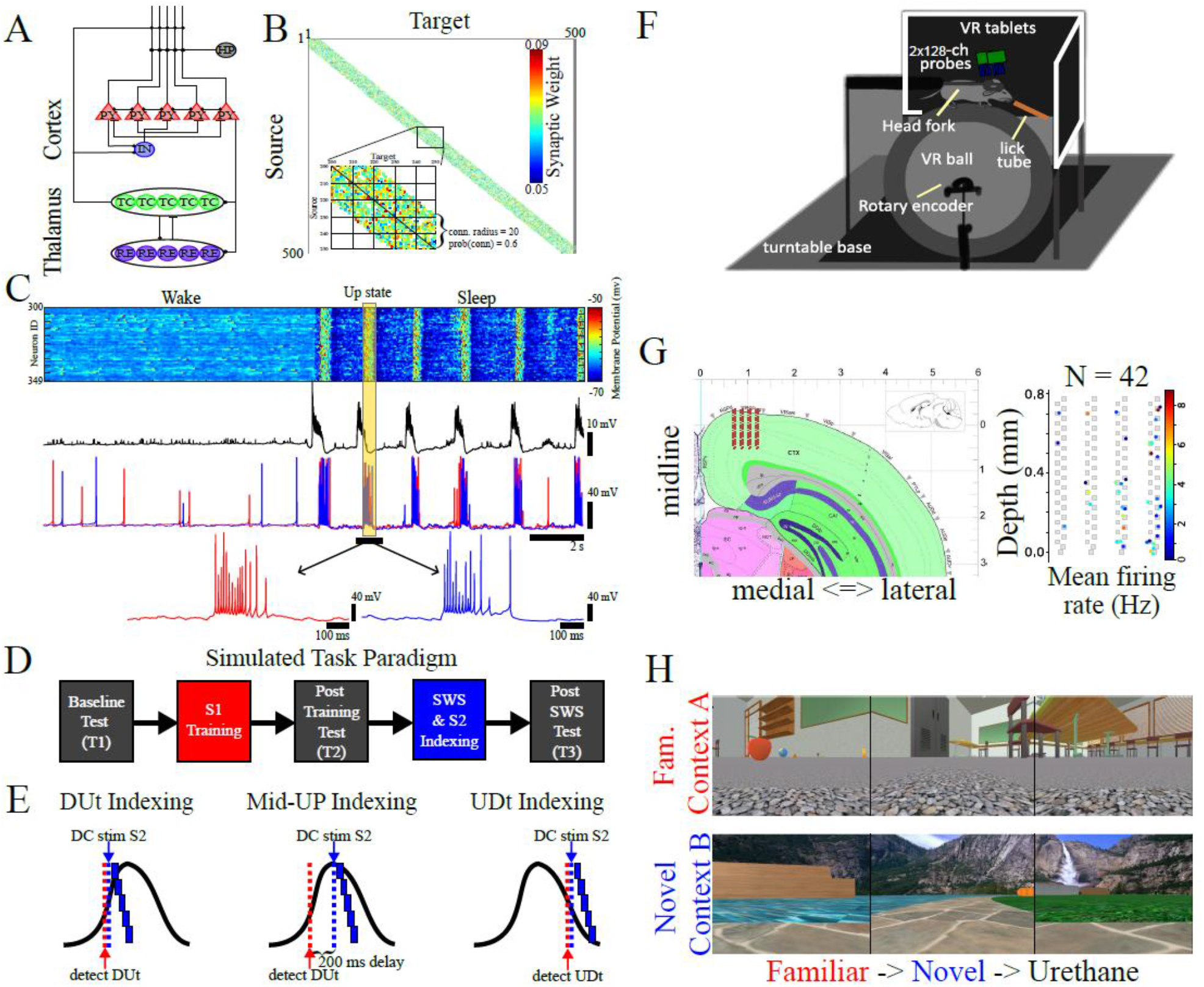
Network model and rodent paradigms. **(A)** Basic network architecture (PY: excitatory pyramidal neurons; IN: inhibitory interneurons; TC: excitatory thalamocortical neurons; RE: inhibitory thalamic reticular neurons). Excitatory (inhibitory) synapses are represented by lines terminating in a dot (bar). **(B)** Initial weighted synaptic matrix for the PYs. The color represents the strength of the AMPA connections between PY neurons, with white indicating the lack of synaptic connection. The inset shows a zoom-in of the subregion where training occurs (PYs 200-249). **(C)** Example of Wake and SWS activity in a subregion (PYs 300-349) of the cortex. Top: The y-axis is the PY index, the x-axis is time, and the color scale indicates membrane potential, with dark red corresponding to spiking activity. Middle: Simulated local field potential (LFP). Bottom: Membrane potential traces of two PYs. One Up state is highlighted. Each trace is expanded below for the 1s interval indicated by the black horizontal line. **(D)** Flow chart of the simulated task paradigm for the network model consisting of memory S1 training (red; i.e. when the “old” memory is embedded into the cortex), SWS and memory S2 indexing (blue; i.e. when input encoding the “new” memory is applied during sleep), and 3 test periods (black; i.e. when pattern completion of the sequence memories is assessed). Each memory is represented by groups of neurons (ten neurons per group) activated in a sequence (e.g., S2=ABCDE). **(E)** Schematic of how hippocampal indexing/SWRs are simulated as external input applied during 3 distinct phases of the cortical Up state during SWS. **(F)** Schematic of behavioral paradigm. Mice were head-fixed over a roller ball with VR tablets displaying a virtual environment. A lick tube dispensed fluid reward, with 4 distinct 128-channel probes recorded neural activity. **(G)** (Left) Coronal section with 128-channel silicon probe map (red square) targeting deep layers of retrosplenial cortex (RSC). (Right) Silicon probe map layout (128 channels, gray squares) with good units (n=42) for one mouse in colored circles overlaid on top. Color corresponds to mean firing rate. **(H)** Animals’ view on three surrounding tablet screens in VR environment for familiar context (context A, top) and novel context (context B, bottom). Experiment involved exposure to a familiar context (red), followed by a novel context (blue), and urethane-induced sleep.

In the basic task paradigm (Figure 1D), a familiar memory (S1) is trained during the awake state. The model then transitions into SWS, during which simulated hippocampal sharp wave–ripple (SWR) input is applied to “index” a novel memory (S2). Testing periods before and after each phase assess potential retroactive and prospective interference. Throughout most of the paper, indexing is applied at one of three cortical Up-state phases (Figure 1E) via online detection: the Down-to-Up transition (DUt), the middle of the Up state (Mid-UP), or the Up-to-Down transition (UDt).

### Rodent paradigm

To test the model predictions *in vivo*, we turned to a familiar/novel context paradigm in rodents that closely matched our simulation paradigm. We analyzed data from ^19^ to determine when familiar vs novel memory traces are reactivated in respect to the phase of the SO. In short (see details in **Methods**), mice were trained to run head-fixed in a 1-D circular track virtual-reality (VR) context for sweetened milk reward at two sites (Figure 1F). Training continued until the mice ran 60 laps at 2 laps/min for three consecutive days. Afterward, craniotomy surgery over retrosplenial cortex (RSC) and secondary motor cortex (M2) was performed, and a catheter was implanted in the upper back to administer urethane in head-fixed mice. The day after surgery (test day), two 128-channel Si probes were used to record single-unit and local-field potentials (LFP) from both brain regions (Figure 1G) ^20^. On the test day, mice ran in a familiar context (context A), followed by a novel context (context B) for either 60 laps or 30 minutes, whichever occurred first. Note that the novel context had a different background texture, object, and reward sites compared to the familiar context to promote orthogonal spatial representation (Figure 1H). After the behavioral session, urethane (subcutaneous 750 mg/kg) was injected to record SWS (total duration: mouse1: 3633.11 s, mouse2: 4398.4 s).

### Memory sequencies

Temporally structured sequences of events are among the most common forms of learned information and are undoubtedly represented in the brain by sequences of neuronal firing. In the model, following ^17,21–23^, each memory was represented as an ordered sequence of activations in cortical neuron populations (e.g., ABC…), where each “letter” denotes a specific neuronal group, and each memory forms a unique “word”. The simulation paradigm - comprising testing, training, and SWS - is illustrated in Figure 2A,B. During wake, the network was trained on memory S1, represented by five sequentially activated cell groups (EDCBA; see Figure 2B, left), serving as a familiar cortical trace. During sleep, hippocampal input encoded a novel memory S2 (ABCDE; see Figure 2B, right). In test sessions, only the first group (E or A) was activated to assess pattern completion (Fig. 2B, middle). Performance was quantified as the distance between the trained template and the network’s response during testing ^17,21^.

**Figure 2.**
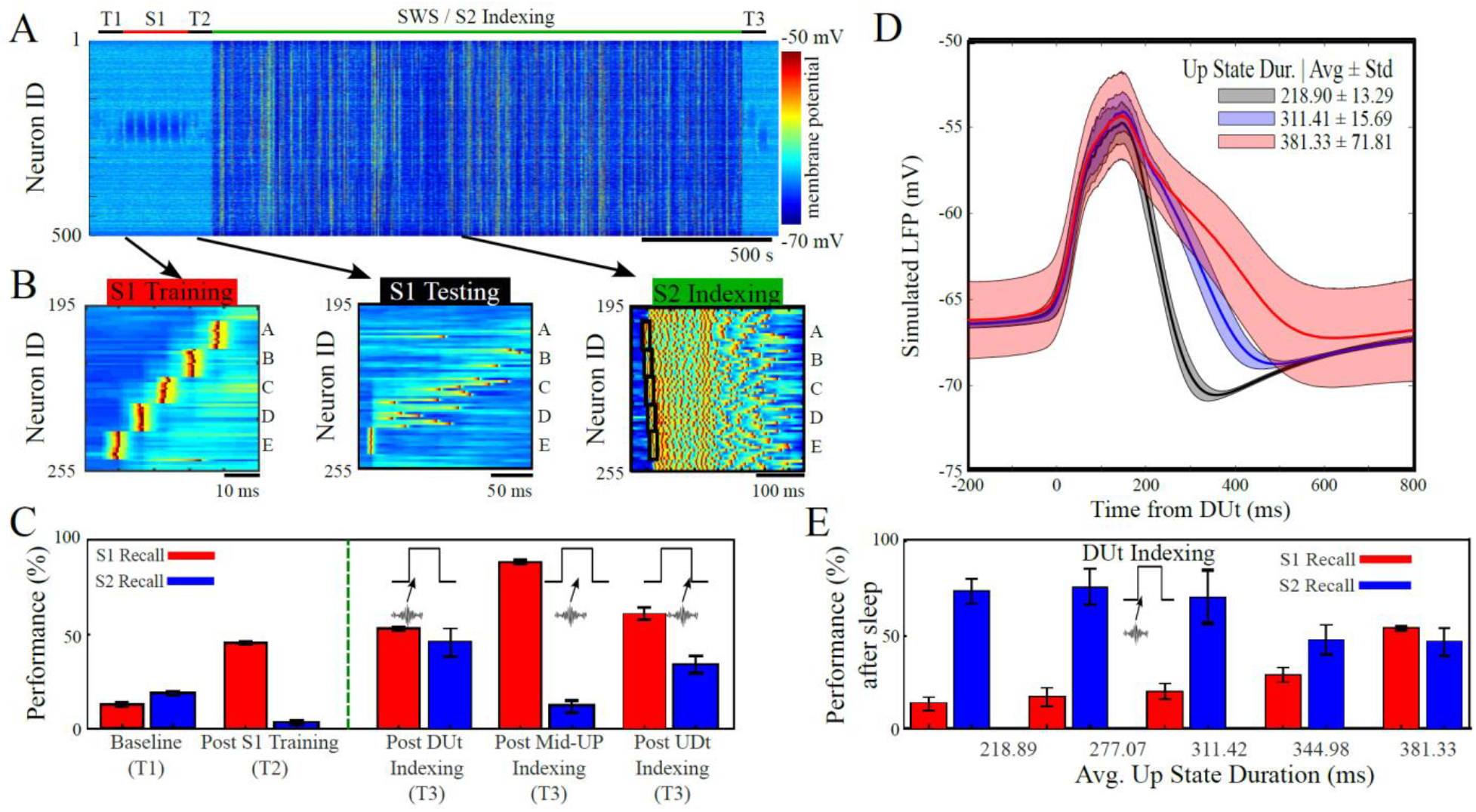
Indexing near the DUt or UDt can drive cortical consolidation of novel memories. **(A)** Network activity during an example simulation depicting membrane potential (color scale) of 500 PYs on the y-axis over time (x-axis). The first memory (familiar, S1) was trained in awake state, that was followed by indexing of the second (novel, S2) memory during SWS. The network undergoes testing periods (T1, T2, and T3; black) at baseline, after S1 training (red), and after SWS / S2 indexing (green). For testing, only the first letter was activated, and performance was evaluated by pattern completion. Examples of network activity during one bout of S1 training (left), during a single test trial (middle), and during a single Up state of SWS when the S2 index (black boxes) was applied near the DUt (right). Bar plots showing familiar S1 (red) and novel S2 (blue) recall performance at baseline (T1), after S1 training (T2), and after SWS (T3). T1 and T2 performance was identical across conditions, while T3 performance (separated by green vertical dashed line) depended on whether the index was applied at the DUt, the Mid-UP, or the UDt. Error bars denote standard deviation. **(D)** Example of average (line) and standard deviation (shading) of simulated LFP traces of the cortical SO, time-locked to the DUt, for 3 distinct sets of simulations with different average Up state durations attained by modifying Ca^2+^-dependent K^+^ current (I_K(Ca)_) in PY neurons. Red - baseline DUt indexing case. **(E)** Bar plots showing recall performance for familiar memory S1 (red) and novel memory S2 (blue) after sleep in simulations with indexing applied at the DUt, across five conditions with varying Up state durations (x-axis). Error bars represent standard deviation. The right-most pair of bars is identical to that shown in panel (C).

### Indexing near Up/Down transitions

We previously reported that when two competing memories were trained sequentially during the awake state, training the second (novel, S2) memory impaired the first (familiar, S1) memory ^17^. This impairment could be reversed by applying SWS after the new training; however, prolonged S2 training led to irrecoverable damage to the familiar S1 memory. This finding suggests that gradual new memory training during wakefulness, along with multiple interleaved episodes of training and subsequent sleep, consistent with procedural learning ^24^, is necessary to enable interference-free learning when cortical synaptic weights are directly modified during awake training.

According to Systems Consolidation Theory ^1,7^, the hippocampus rapidly encodes new declarative memories for later consolidation in the cortex during SWS. A key element of this process is cortical replay triggered by hippocampal SWRs ^8^ (i.e. “hippocampal indexing”). Here, we asked how replay of novel hippocampus-dependent memories can avoid overwriting familiar memory traces already encoded in overlapping cortical neurons populations, thereby enabling interference-free encoding of new memories into the cortical network.

To model consolidation of a new memory, the cortical network - previously trained on sequence S1 (EDCBA) - was subjected to “hippocampal indexing”, simulating hippocampal replay of a novel memory S2 (Figure 2A). To maximize the potential for interference under our STDP rules, S2 was designed as the reverse of S1 (i.e., S2: ABCDE). Indexing was implemented by detecting a specific phase of the Up state and delivering a sequence of suprathreshold DC pulses to activate the S2 sequence (Figure 2B, right). Specifically, we compared three indexing conditions: the Down-to-Up transition (DUt), the middle of the Up state (Mid-UP), or the Up-to-Down transition (UDt). DUt or UDt phases were detected online using the global LFP (see Methods for details), and indexing was triggered immediately upon detection. For the Mid-UP condition, indexing was applied with a 200 ms delay after DUt detection. Notably, Up states in the model had an average duration of 400 ms. Memory performance was assessed in the awake state at three time points: baseline (T1), after S1 training (T2), and after SWS with S2 indexing (T3), across all three indexing conditions (Figure 2C).

Familiar memory S1 performance improved following initial awake training (Figure 2C; T2, red) and further increased after sleep with S2 indexing (T3, red), across all three conditions. In contrast, S2 performance after sleep depended on the timing of hippocampal indexing (T3, blue). With DUt indexing (T3, left), recall improved for both S1 and S2, yielding comparable performance. Mid-UP indexing (T3; middle) supported consolidation of familiar S1 memory but failed to consolidate the novel S2 memory. With UDt indexing (T3, right), recall again improved for both memories, but the network was biased toward S1 over S2. Thus, indexing near Up or Down state transitions supported consolidation of both familiar and novel memories, while Mid-UP indexing failed to consolidate the novel memory.

To ensure that our findings were not specific to a particular memory representation, we reduced interference between the two memories by modifying the sequences from fully to partially overlapping (Suppl. Figure 1A) and observed qualitatively similar results (Suppl. Figure 1B–D). These findings were further confirmed using more complex training protocols, in which the familiar memory (S1) was first consolidated during one sleep phase, and the novel memory (S2) was encoded during a subsequent, separate sleep phase (Suppl. Figure 2).

Since Mid-UP indexing did not support consolidation of the novel memory (S2) but further enhanced performance of the familiar memory (S1), we hypothesized that this phase of the Up state plays a role in protecting S1 from interference by S2. To test this, we ran simulations using DUt indexing while reducing the Up-state duration by increasing the conductance of Ca^2+^-dependent K^+^ currents in cortical PYs - one of the mechanisms involved in Up-state termination ^25,26^ (Figure 2D). We found that even the shortest Up states could support consolidation of the novel memory S2 (Figure 2E; blue bars), but shorter Up-state durations substantially reduced S1 performance (Figure 2E; red bars). These results support our hypothesis that the complementary learning systems model does not inherently protect old memories, which can still be disrupted if new memory indexing targets overlapping neuron and synapse populations.

### Successful indexing induces interleaved replay

To understand how the sleep model with typical length Up states (i.e. long; ~ 400s) can consolidate novel memories without disrupting familiar ones, we analyzed synaptic reactivation using approach previously applied in ^17,27^. Since indexing a new memory is expected to promote its reactivation, we examined reactivation separately during the Indexed Phase (i.e., when the indexing signal was applied) and the Non-Indexed Phase(s) (i.e., the remainder of the Up state; see examples in Figure 3A–C, right). Briefly (see **Methods** for details), for each synapse in the training region, we counted the number of Up states in which it showed net long-term potentiation (LTP), separately for the Indexed and Non-Indexed phases (Figure 3A–C, left vs middle plots). Next, due to the model’s geometry, each synapse was classified as supporting either S1 (if projecting from high to low cell indexes) or S2 (if projecting from low to high). Thus, e.g., net LTP in synapses supporting S1 (red boxes in Figure 3A–C) indicates reactivation of the S1 memory at a given Up state.

**Figure 3.**
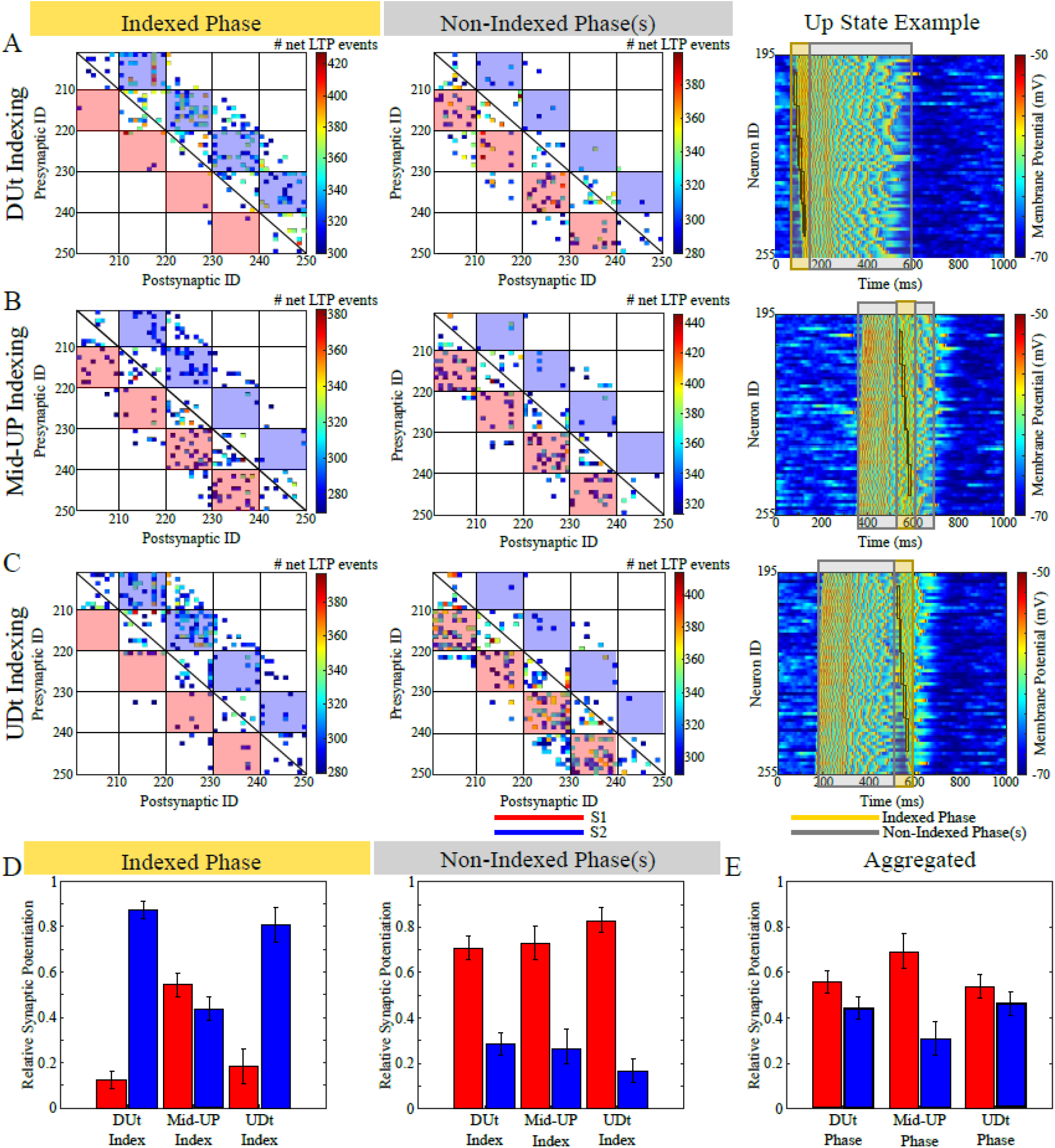
Phase-specific memory replay during cortical Up states. **(A-C)** Heatmaps show the number of Up states (~450 total per simulation) in which each synapse exhibited net LTP. These events were counted separately during the Indexed Phase - when the artificial index was applied (left panels) - and the Non-Indexed Phase(s), i.e., periods without indexing within each Up state (middle panels). In three independent conditions, the index was applied at different times within the Up state: (A) DUt, (B) Mid-Up, and (C) UDt. Red and blue backgrounds indicate synapses contributing to recall of S1 and S2, respectively, based on connection directionality. To avoid visual clutter, only the top 30% of synapses are shown. The Up state example (right) shows a single Up state for each of the three indexing conditions, depicting the indexed phase (yellow shading), the non-indexed phases (gray shading), and the applied index (black boxes). **(D)** Bar plots show the relative amount of synaptic potentiation for S1 (red) versus S2 (blue) during the Indexed Phase (left) and Non-Indexed Phase(s) (right). Note S1 was preferentially reactivated during Non-Indexed phase, whereas S2 was reactivated during the Indexed phase when the index was applied near DUt or UDt. **(E)** Bar plots show relative synaptic potentiation during the DUt, Mid-Up, and UDt phases, with data aggregated across all three indexing conditions under the assumption of equal likelihood. Error bars in (D) and (E) indicate standard deviation.

For DUt indexing (Figure 3A), we observed robust reactivation of the novel S2 memory (blue boxes) during the Indexed Phase and of the familiar S1 memory (red boxes) during the Non-Indexed Phase. With Mid-UP indexing (Figure 3B), only S1 was strongly reactivated across both phases. Similar to DUt, UDt indexing (Figure 3C) yielded strong S2 reactivation during the Indexed Phase and S1 reactivation during the Non-Indexed Phase. Notably, Mid-UP indexing triggered a modest initial reactivation of S2 (A→B) but failed to drive full sequence replay, suggesting that ongoing activity in the middle of an UP state is less susceptible to external input.

This data is summarized in Figure 3D. We summed the LTP event counts within the red and blue boxes (corresponding to S1 and S2, respectively) and normalized by the total number of LTP events.

Familiar memory (S1; red) was robustly reactivated throughout the Non-Indexed Phases across all indexing conditions (Figure 3D, right). In contrast, reliable reactivation of the new memory (S2; blue) occurred only when indexing was timed near Up/Down transitions (DUt or UDt) (Figure 3D, left). For Mid-UP indexing, the network instead spontaneously reactivated S1 during the Indexed Phase, despite attempts to index S2 (see Figure 3D, left – Mid-UP Index). This result also explains why DUt indexing during SWS with very short Up states led to damage of old memories (Figure 2G): the network lacked sufficient time to reactivate S1 and protect it from interference.

Because the rodent paradigm did not permit recording SWR timing (see below), we aggregated data across all three indexing conditions in Figure 3E (see Methods), assuming ripples occur equally across all Up-state phases. We then plotted relative synaptic reactivation during DUt, Mid-Up, and UDt phases. As expected, reactivation of familiar S1 and novel S2 was comparable during DUt and UDt, since in the absence of indexing the network reactivates S1, whereas indexing drives S2. Overall, the model predicts that when Up states are sufficiently long, familiar memory reactivation dominates the middle phase, while reactivation near Up–Down transitions can reflect either familiar or novel memories, depending on SWR timing. Thus, familiar and novel memory traces replay in an interleaved manner within individual Up states, protecting familiar memories from interference by hippocampal indexing of novel ones.

We repeated this analysis for net LTD events (Suppl. Figure 3). For Mid-UP and UDt indexing, the results were essentially the inverse of the net LTP findings, consistent with the anti-symmetric STDP kernel used during sleep. For DUt indexing, the outcome was unexpected. First, unlike LTP, the relative amount of synaptic depression was similar for both memories across the Up state (Suppl. Figure 3A and 3D). Second, the number of net LTD events was much lower than LTP across both phases of the Up state (compare color scales in Figure 3A and Suppl. Figure 3A). Given that STDP rule is anti-symmetric, this suggests that DUt indexing may preferentially strengthen unidirectional synapses (i.e., those not part of bidirectionally-connected pairs on neurons). These results imply that DUt and UDt indexing promote distinct reactivation dynamics and may differentially shape systems consolidation by favoring unidirectional (DUt) or bidirectional (UDt) synaptic configurations.

### Rodent model recapitulates familiar/novel memory reactivation dynamics

Due to our interest in spatially selective firing and the large fraction of reward-selective firing in M2 in the dataset, we focused our analysis on RSC data (Figure 1C) and excluded M2 units (Suppl. Figure 4A-D). After detecting NREM and REM epochs, we calculated the Up and Down states (Suppl. Figure 4E-H). Principal-component analysis (PCA) based reactivation analysis ^8,28^ was applied to Up states to estimate the strength of novel and familiar context reactivation during sleep, using population activity from behavior sessions in individual contexts as templates. We focused on Up states with a duration > 300 ms to better align with modeling work. Each Up state was divided into three equal parts, and the probability of novel context reactivation being greater than familiar context reactivation was estimated for each part in 4-second windows (Figure 4A). This provided three distributions indicating the probability of significant novel context reactivation and familiar context reactivation (1 – [probability of novel]) every 4 seconds. We observed a higher novel context reactivation in the last third followed by the first third, and finally the middle third of the Up-states for both animals (Figure 4B).

**Figure 4.**
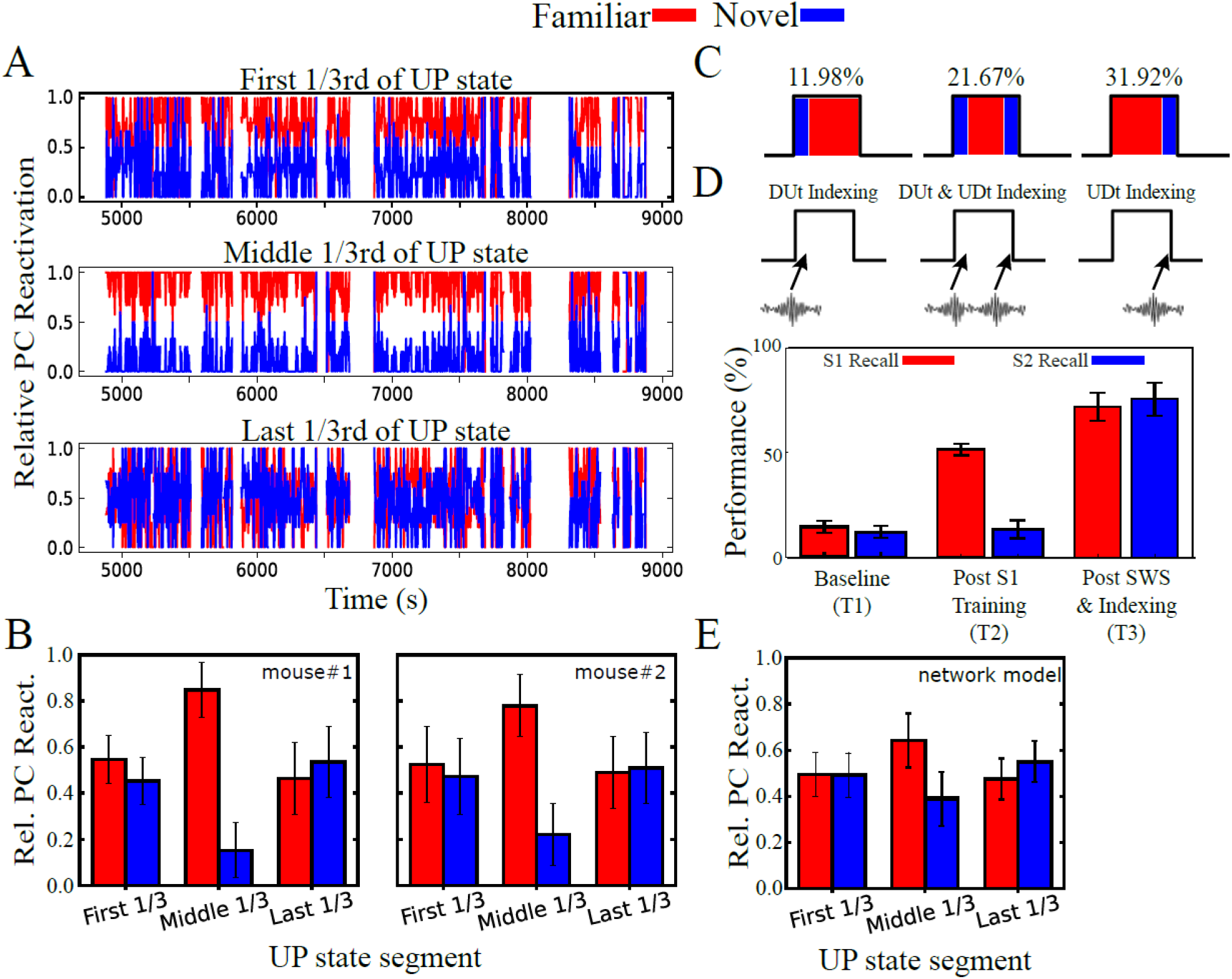
Familiar memory replay dominates the middle of Up states and mixes with novel memory replay near UDt and DUt transitions in rodents. **(A)** Example relative PC reactivation for novel and familiar context reactivation in 4 s windows for three equal parts (shown in each row) of individual Up states. Note that both the last (third row) and first 1/3rd (top row) show higher novel context (blue) reactivation probability than the middle 1/3rd (second row). **(B)** Mean ± sem familiar (red) and novel (blue) context relative PC reactivation for each 1/3rd segment of UP-states for both animals. (**C**) Schematic illustrating dominant replay dynamics of familiar (red) and novel (blue) memories found empirically during cortical Up states. 1. Novel memory replay at the DUt with familiar memory replay dominating the rest of the Up state (top); 2. Novel memory replay at both the DUt and UDt with familiar replay dominating in the Mid-UP state (middle); and 3. Familiar replay dominating the DUt and Mid-UP and novel replay at the UDt (bottom). **(D)** Schematic depicting how indexing was simulated in a network model where the type and frequency of indexing was applied stochastically to match the empirical probabilities obtained in (C). See Suppl. Figure 5 for more details. **(E)** Bar plots showing S1 (red) and S2 (blue) recall performance at baseline (T1), after S1 training (T2), and after SWS (T3); error bars denote standard deviation. (**E**) Mean ± sem of relative PC reactivation for the familiar (S1; red) and novel (S2; blue) memory sequences in the network model.

Changing analysis parameters (Up state window size) did not alter our qualitative results (Suppl. Figure 4I). These findings are remarkably in line with the predictions of the model shown in Figure 3D and 3E (see below).

Although we cannot say empirically whether SWRs are responsible for the novel memory replay, we were able to summarize the *in vivo* observations into three patterns of replay during Up states, along with the frequency of occurrence (Figure 4C). We then ran a model simulation in which we indexed the novel memory stochastically on each Up state at either the DUt, UDt, or both (Figure 4D; top), in accordance with empirical observations. Note that for this simulation, we used partially overlapping representations for the memory sequences (see Suppl. Figure 1) rather than fully overlapping representations as this is more likely to be the case *in vivo*. Following SWS, recall performance was maintained and even increased for the familiar memory, while the novel one was robustly consolidated (Figure 4D; bottom). These results are in very good agreement with the in vivo data shown in Figure 4B. Remarkably, this simulation suggests that combining DUt and UDt indexing in this manner results in a synergy, as it out-performed all simulations where DUt or UDt indexing were applied alone (compare to Suppl. Figure 1B,D). Finally, PCA-based reactivation analysis, as used for rodent data, was applied to the model with stochastic indexing (Figure 4E), once again showing good qualitative agreement between simulated and in vivo data (see also Suppl. Figure 5).

## DISCUSSION

Humans and animals possess a profound capability to continuously adapt throughout their lifespans, aided by their brain’s immense plasticity. The brain can learn rules and patterns from relatively few examples, incorporate new facts into its corpus of knowledge, and generalize beyond its experienced sensorium to plan, hypothesize, and imagine. While much deserved attention has been paid to how these learning processes proceed as animals acquire new experiences while awake, a vast interdisciplinary literature highlights the critical role that sleep has in these functions through a process termed memory consolidation^1–5^.

Despite extensive characterization of the behavioral and cognitive functions facilitated by sleep ^29–33^, our understanding of memory consolidation at the neuronal level is largely limited to two fundamental ideas: (1) neuronal ensembles representing recent memories are reactivated during sleep ^34,35^, and (2) declarative memories are initially learned by the hippocampus and later transferred to the cortex during slow-wave sleep (SWS) ^1,7^ – Systems Consolidation Theory ^36–38^, also known as Complementary Learning Systems (CLS) Theory ^1,39^. Particular attention has been paid to the replay which occurs during sharp wave-ripples (SWR) in the hippocampus ^40,41^ and Up states of the slow oscillation (SO) in the cortex ^8,42–44^. This powerful idea, proposed in the 1990s ^1^, remains central to our understanding of declarative memory consolidation. However, it does not guarantee interference-free learning - new memories can still disrupt familiar ones during consolidation phase, as we report here.

In this new study, we propose an important extension to CLS, named Structured Cortical Replay (SCoRe), to explain how new memories encoded by the hippocampus and carried over by hippocampal ripples avoid overwriting cortical traces of familiar, overlapping memories already encoded by the cortex. SCoRe posits that reactivation of both new (triggered by hippocampal SWRs) and old (previously encoded in cortical weights) memory traces is tightly coordinated - interleaved - within individual Up states of slow-wave sleep, such that novel memories tend to replay at the beginning and end of the Up state, while familiar memories tend to replay in the middle. This hypothesis, predicted by the model, was confirmed in *in vivo* experiments in which animals were first trained in environment 1 and later switched to environment 2, followed by sleep.

While we do not have access to SWR occurrence times in this study, our predictions qualitatively align with recent observations showing that ripples primarily occur around the DUt and UDt phases of the SO in RSC ^45^. Although there is a slight quantitative discrepancy - empirical data show ripple peaks occurring later than the ideal timing predicted by our model - we suspect this may be due to the one-dimensional nature of our model, which could influence the time course of transitions between Up and Down states. Interestingly, another recent study ^46^ suggested that replays of older and newer memories within the hippocampus are also interleaved to avoid interference. While our study focuses on cortical replay and interference between newer hippocampus-dependent and older hippocampus-independent (purely cortical) memories, it points to a similar overarching strategy - interleaving reactivations to mitigate interference.

Our *in vivo* data revealed three distinct interleaved replay patterns: familiar memory was consistently replayed in the middle of the Up state, while novel memory was replayed at the DUt (11.98%), UDt (31.92%), or both phases (21.67%). Intriguingly, our model predicts that DUt and UDt indexing have distinct effects at the synaptic level, and that combining them yields a synergistic effect - producing greater consolidation gains than either alone. It will be important to further investigate the functional roles of these two ripple classes. The SCoRe enables the formation of novel cortical memory traces while minimizing interference and damage to familiar ones, likely relying on bidirectional cortico-hippocampal dialogue^12–15^. Another important advantage is that replaying the novel memory near the beginning of an Up state likely triggers replay of overlapping familiar memories later in the same Up state, thereby enabling similarity-weighted interleaved learning (SWIL) ^39,47^.

Our study makes several important empirical predictions that could be tested in future experiments. These include: 1) short-duration Up states coupled to a SWR preferentially replay novel memories in the cortex. This could be empirically assessed with a behavioral paradigm similar to ours but with paired CA1 recordings to determine SWR timing. Furthermore, the duration of Up states can be pharmacologically manipulated using bicuculline methiodide to progressively suppress fast-acting inhibition ^48^, and we predict this would lead to progressive overwriting of familiar cortical memories. 2) Disrupting SWRs during the UDt and DUt will impair consolidation of novel memories, which could be assessed through online detection of SWRs and SOs and selective disruption of SWRs occurring at these two phases. 3) Our net LTP/LTD analyses suggest that SWRs arriving at the UDt cause greater interference to overlapping familiar memories than those arriving at the DUt. This could potentially be assessed using a closed-loop auditory stimulation paradigm to trigger SWRs at the appropriate phase of the cortical Up state during sleep ^49^, although the reactivation content of the SWRs would need to be accounted for. 4) Our stochastic indexing simulation found that a random combination of DUt and UDt indexing outperforms either alone. This could be tested by using optogenetic inhibition to suppress SWRs during the DUt or the UDt and comparing the resulting behavioral performance on both tasks to natural sleep.

From a neuronal ensemble perspective, coordinated replay of familiar and novel memories effectively segregates synaptic resources between tasks. Indeed, we previously showed using biophysical spiking models that synaptic reactivation during sleep can rescue memories previously damaged by sequential training ^17,22,23,50^. For overlapping memories, interleaved training replaces the original single attractor encoding the familiar memory with multiple attractors encoding both familiar and novel memories - embedded within the same neuronal and synaptic population ^17,50^.

These ideas were further extended to ANNs, where we found that sleep-like replay restored performance across multiple tasks trained sequentially - Sleep Replay Consolidation (SRC) ^51^. The SRC approach, while effective, has the drawback that new training first damages old memories, which are then recovered during a subsequent sleep-like phase; therefore, the initial damage must not be too severe. SRC is likely analogous to how the brain learns procedural tasks (e.g., skills like playing tennis). Procedural learning is typically slow and involves multiple training episodes interleaved with sleep cycles. This gradual process allows normal night sleep to “repair” damage to old memories and converge toward a stable joint representation ^50^ over the course of multiple nights and training episodes.

Our new study proposes a novel strategy for ANNs that may more closely mimic declarative memory learning. This would require implementing a dual memory system, in which one (hippocampal, ANN_H_) network rapidly acquires a new task and reactivates it during a sleep-like phase (possibly in spiking mode, following the strategy from ^51^), projecting reactivation patterns to a second (cortical, ANN_C_) network that concurrently reactivates relevant existing memory traces. This mechanism could allow new memory traces to be embedded in ANN_C_ without causing significant disruption to preexisting ones.

The sleep-based approach for continual learning offers advantages over existing methods for mitigating catastrophic forgetting, which fall into two broad categories (reviewed in ^4,52^): rehearsal and pseudorehearsal ^1,9–11^, i.e., storing or generating old data samples to mix with new ones during training; and regularization-based methods ^53–56^, which modify plasticity rules by adding constraints to gradient descent that preserve weights important for previously learned tasks. Rehearsal requires storing increasingly large amounts of data or building increasingly complex models to generate representative samples, which is impractical in lifelong learning scenarios. Regularization, on the other hand, becomes less effective when tasks are highly overlapping or when new learning requires modification of weights previously deemed important, limiting flexibility and scalability over long task sequences.

In sum, our study reveals fundamental principles of how the brain leverages the dual-memory (cortico-hippocampal) system for fast, efficient, and interference-free continual learning. Beyond neuroscience, it may also inform the development of more robust continual learning solutions in AI.

## METHODS

### Thalamocortical model methods

All simulations were carried out using custom C++ code and all analyses were performed with standard MATLAB and Python functions. Data are presented as mean standard error of the mean (SEM) unless otherwise stated. For each condition a total of 6 simulations with different random seeds were used for statistical analysis.

### Network architecture

Throughout this study, we make use of a slightly modified version of a thalamocortical network which has been previously described in detail ^17,27^. In brief, the network consisted of a cortical module containing 500 excitatory pyramidal neurons (PYs) and 100 inhibitory interneurons (INs), and a thalamic module containing 100 excitatory thalamocortical neurons (TCs) and 100 inhibitory reticular interneurons (REs). Connectivity in the network was determined by cell type and a local radius (see Figure 1), and excitatory synapses were mediated by AMPA and/or NMDA currents, while inhibitory synapses were mediated by GABA_A_ and/or GABA_B_ currents.

In the cortex, PYs synapsed onto PYs and INs with a radii of R_AMPA(PY-PY)_ = 20, R_NMDA(PY-PY)_ = 5, R_AMPA(PY-IN)_ = 1, and R_NMDA(PY-IN)_ = 1. All connections were deterministic within these radii, expect for AMPA synapses between PYs, which had a 60% probability of connection. Additionally, INs synapsed onto PYs with a radius of R_GABA-A(IN-PY)_ = 5. In the thalamus, TCs synapsed onto REs with a radius of R_AMPA(TC-RE)_ = 8 and REs synapsed onto REs and TCs with radii of R_GABA-A(RE-RE)_ = 5, R_GABA-A(RE-TC)_ = 8, and R_GABA-B(RE-TC)_ = 8. Between the cortex and thalamus, TCs synapsed onto PYs and INs with radii of R_AMPA(TC-PY)_ = 15, R_AMPA(TC-IN)_ = 3, while PYs synapsed onto TCs and REs with radii of R_AMPA(PY-TC)_ = 10, and R_AMPA(PY-RE)_ = 8.

### Wake – Sleep transitions

To model the state transitions between awake and N3 sleep, we modulated the intrinsic and synaptic currents of our neuron models to account for differing concentrations of neuromodulators that partially govern these arousal state transitions. As these mechanisms have been described in detail in ^16^, here we will simply outline the approach. The model included the effects of changing acetylcholine (ACh), histamine (HA), and GABA concentrations as follows: ACh – by modulating the potassium leak current in all cell types, as well as excitatory AMPA synapses within the cortex; HA – by modulating the hyperpolarization-activated cation current in TC cells; and GABA – by modulating inhibitory GABAergic synapses within the cortex and thalamus. To transition the network from awake to sleep, we modeled the effects of reduced ACh and HA but increased GABA concentrations to reflect experimental observations ^57^.

### Intrinsic currents

All cell types were modeled using the Hodgkin-Huxley formalism, and cortical PYs and INs contained dendritic and axo-somatic compartments that have been previously described ^22^. The dynamics of the membrane potential were modeled according to:

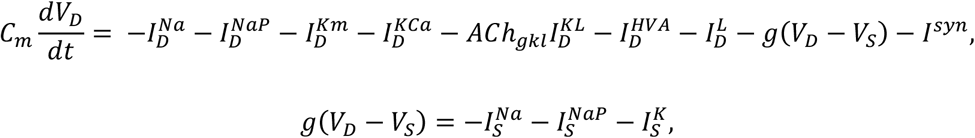

where *C*_*m*_ is the membrane capacitance, *V*_*D*_,_*S*_ are the dendritic and axo-somatic membrane voltages respectively, *I*^*Na*^ is the fast sodium (Na^+^) current, *I*^*NaP*^ is the persistent Na^+^ current, *I*^*Km*^ is the slow voltage-dependent non-inactivating potassium (K^+^) current, *I*^*KCa*^ is the slow calcium (Ca^2+^)-dependent K^+^ current, *ACh*_*gkl*_ represents the change in K^+^ leak current *I*^*KL*^ which is dependent on the level of ACh during the different arousal states, *I*^*HVA*^ is the high-threshold Ca^2+^ current, *I*^*L*^ is the chloride (Cl^−^) leak current, *g* is the conductance between the dendritic and axo-somatic compartments, and *I*^*syn*^ is the total synaptic current input to the neuron. IN neurons contained all intrinsic currents present in PY with the exception of the *I*^*NaP*^. All intrinsic ionic currents (*I*^*j*^) were modeled in a similar form:

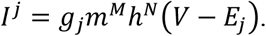

where *g*_*j*_ is the maximum conductance, *m* (activation) and *h* (inactivation) are the gating variables, *V* is the voltage of the compartment, and *E*_*j*_ is the reversal potential of the ionic current. The gating variable dynamics are described as follows:

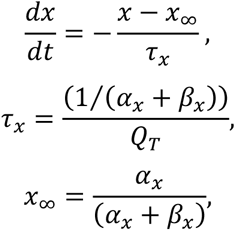

where *x* = *m* or *h, τ* is the time constant, *Q*_*T*_ is the temperature related term, *Q*_*T*_ = *Q*^((*T*™23)/10)^ = 2.9529, with *Q* = 2.3 and *T* = 36.

In the thalamus, TCs and REs contained a single compartment with membrane potential dynamics given by:

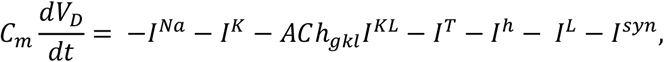

where *I*^*Na*^ is the fast Na^+^ current, *I*^*K*^ is the fast K^+^ current, *I*^*KL*^ is the K^+^ leak current, *I*^*T*^ is the low-threshold Ca^2+^ current, *I*^*h*^ is the hyperpolarization-activated mixed cation current, *I*^*L*^ is the Cl^−^ leak current, and *I*^*syn*^ is the total synaptic current input to the neurons. The *I*^*h*^ current was only expressed in TCs. The influence of histamine (HA) on *I*^*h*^ was implemented as a shift in the activation curve by *HA*_*gh*_ as described by:

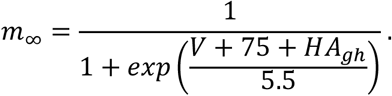

### Synaptic currents

The equations for our synaptic current models have been described in detail in our previous studies ^16,58^. To model the effects of ACh and GABA, we modified the standard equations as follows:

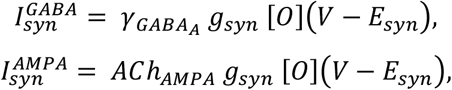

where *g*_*syn*_ is the maximal conductance at the synapse,[*O*] is the fraction of open channels, and *E*_*syn*_ is the channel reversal potential (E_GABA-A_ = −70 mV, E_AMPA_ = 0 mV, and E_NMDA_ = 0 mv). The parameter 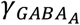 modulated the GABA synaptic currents for IN-PY, RE-RE, and RE-TC connections. For Ins 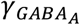 was 0.22 and 0.44 for awake and N3 sleep, respectively, while for 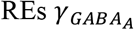 was 0.6 and 1.2. *ACh*_*AMPA*_ defined the influence of ACh levels on AMPA synaptic currents for PY-PY, TC-PY, and TC-IN. For PYs *ACh*_*AMPA*_ was 0.133 and 0.4332 for awake and N3 sleep, respectively, while for TCs *ACh*_*AMPA*_ was 0.6 and 1.2.

In addition to spike-triggered post-synaptic potentials (PSPs), spontaneous miniature PSPs (mPSPs) were implemented for both excitatory and inhibitory synapses within the cortex. The dynamics are similar to the typical PSPs described above, but the arrival times were governed by an inhomogeneous Poisson process where the next release time *t*_*release*_ is given by:

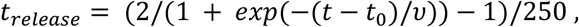

where *t*_0_ is the time of the last presynaptic spike, and *υ* was the mPSP 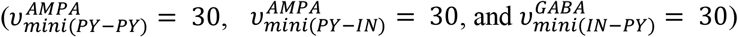. The maximum conductances for mPSPs were 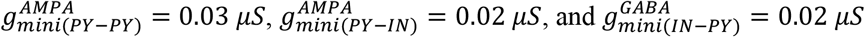.

Finally, short-term synaptic depression was also implemented in AMPA synapses within the cortex. To model this phenomenon, the maximum synaptic conductance was multiplied by a depression variable (*D* ≤ 1), which represents the amount of available “synaptic resources” as described in ^26^. This short-term depression was modeled as follows:

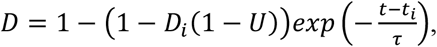

where *D*_*i*_ is the value of *D* immediately before the *i*_*th*_ event, (*t* − *t*_*i*_) is the time after the *i* _*th*_ event, *U* = 0.073 is the fraction of synaptic resources used per action potential, and *τ* = 700*ms* is time constant of recovery of synaptic resources.

### Spike-timing-dependent plasticity

The potentiation and depression of AMPA synapses between PYs were governed by the following spike-timing-dependent plasticity (STDP) rule:

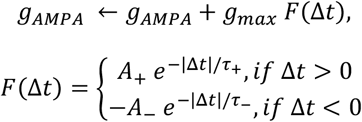

where *g*_*max*_ was the maximal conductance of *g*_*AMPA*_, *F* was the STDP kernel, and □ *t* was the relative timing of the pre- and post-synaptic spikes. The maximum potentiation/depression were set to A_+/-_ = 0.002, while the time constants were set to t_+/-_ = 20 ms. A_-_ was reduced to 0.001 during training to reflect the effects of changes in acetylcholine concentration during focused attention on synaptic depression during task learning observed experimentally ^59–61^.

### Sequence training and testing

Training and testing of memory sequences was performed similarly to our previous study ^22^. In brief, each sequence was comprised of the same 5 groups of 10 PYs (i.e PYs 200 - 249), with Sequence 1 (S1) ordered E(240-249), D(230-239), C(220-229), B(210-119), A(200-209), and Sequence 2 (S2) ordered A(200-209), B(210-219), C(220-229), D(230-239), E(240-249). Each training bout consisted of sequentially activating each group via a 10 ms direct current pulse with a 5 ms delay between group activations. Training bouts occurred every 1 s during the training period. This training structure was chosen to ensure strong interference between S1 and S2 according to our STDP rule. Test bouts occurred every 1 ms during testing periods, in which only the first group in each sequence was activated (E for S1; A for S2), and recall performance was measured based on the extent of pattern completion for the remainder of the sequence within a 350 ms window.

### Sequence performance measure

A detailed description of the performance measure used during testing can be found in ^22^ and the code is available in (https://github.com/o2gonzalez/sequencePerformanceAnalysis)^62^. Briefly, the performance of the network on recalling a given sequence following activation of the first group of that sequence was measured by the percent of successful sequence recalls. We first detected all spikes within the predefined 350 ms time window for all 5 groups of neurons in a sequence. The firing rate of each group was then smoothed by convolving the average instantaneous firing rate of the group’s 10 neurons with a Gaussian kernel with window size of 50 ms. We then sorted the peaks of the smoothed firing rates during the 350 ms window to determine the ordering of group activations. Next, we applied a string match (SM) method to determine the similarity between the detected sequences and an ideal sequence (ie. A-B-C-D-E for S1). SM was calculated using the following equation:

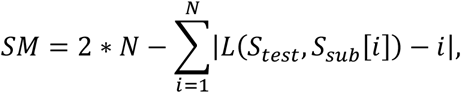

where *N* is the sequence length of *S*_*test*_, *S*_*test*_ is the test sequence generated by the network during testing, *S*_*sub*_ is a subset of the ideal sequence that only contains the same elements of *S*_*test*_, and *L*(*S*_*test*_, *S*_*sub*_[*i*]) is the location of the element *S*_*sub*_[*i*] in sequence *S*_*test*_. *SM* was then normalized by double the length of the ideal sequence. Finally, the performance was calculated as the percent of recalled sequences with *SM* ≥ *Th* = 0.8, where *Th* is a threshold indicating that the recalled sequence must be at least 80% similar to the ideal sequence to be counted as a successful recall as previously done in ^22^.

### Online DUt and UDt detection

Online detection of the DUt and UDt during the simulations was conducted by tracking the simulated global LFP in the same manner as reported previously ^23^. Briefly, the LFP was approximated by calculating mean membrane potentials of all the cortical excitatory neurons, and it had a bimodal distribution during N3 sleep, where one peak corresponded to the Up state and another peak to the Down state. The trough of the distribution was selected as a threshold to separate Up and Down state. The onset of Up or Down state was then defined as the moment when LFP value crossed the threshold. For Mid-UP indexing, the onset of the Up state was detected, followed by a 200ms delay before the simulated index was applied.

### Synaptic potentiation/depression analysis

Synaptic potentiation/depression was computed by utilizing an offline implementation of STDP. For the sake of brevity, we describe this analysis for the DUt indexing case, with the Mid-UP and UDt indexing cases following similarly. First, each Up state was identified by detecting the DUt and UDt times during SWS for a given simulation. Each Up state was then divided into an Indexed Phase which began at the application of the index (i.e. the DUt) and lasted for 80 ms, and a Non-Indexed Phase which began at the end of the Indexed Phase and lasted until the end of the Up state (i.e. the UDt).

Next, focusing on a single Up state and a single Phase (e.g. Indexed Phase), the PY spike times were inserted into an offline STDP implementation which was identical to the one used online in simulation. Using this implementation, for a given pair of synaptically connected PYs, we computed all LTP and LTD events which occurred during this phase of the Up state and then summed these events. If the sum was positive/negative, we considered this a net LTP/LTD event for that pair of PYs during a particular phase of a given Up state. This was then repeated for each Up state, with the total number of net LTP/LTD events being recorded for each pair of PYs. The same was then done for the opposite Phase (i.e. Non-Indexed Phase).

The aggregated potentiation plots (Figure 3E) were generated as follows: let 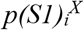 represent the relative synaptic potentiation for S1 during phase X ∈ {Indexed, Non-Indexed} with indexing condition *i* ∈ {DUt, Mid-UP, UDt}. Also let *P(S1)*_*i*_ represent the aggregated relative synaptic potentiation for S1 during the phase *i* ∈ {DUt, Mid-UP, UDt}. Then

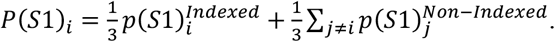

The same follows for *P(S2)*_*i*_, as well as for synaptic depression.

### In vivo methods

#### Animals and Surgical Procedures

Methods were previously described in Kilianski, 2021 ^19^. Briefly, experimental protocols were approved by the Institutional Animal Care and Use Committee of the University of California, Irvine. Four 12-week-old C57BL/6J mice were obtained from Jackson Labs. After headbar surgery, mice were housed individually on a 12-hour on/off light cycle and received limited daily food to maintain their body weight between 75%-80% of their pre-surgery *ad libitum* weight. Mice were first implanted with stainless steel headbars on the skull. After meeting the behavior performance criteria (described below), craniotomy was performed over the secondary motor cortex (M2, AP + 2.5 mm, ML - 1 mm) and retrosplenial cortex (RSC, AP - 3mm, ML – 0.75 mm). A 0.4 mm diameter catheter was implanted under the mice’s upper back for subcutaneous urethane injections.

### Behavior protocol: training and test

After headbar surgery recovery, mice were trained to run for sweetened milk reward in a head-fixed visual virtual reality (VR). They ran on a suspended styrofoam ball, and a rotary encoder was used to update the position on the 1-D circular track VR context (familiar context A) displayed on three tablets (front, and 90° to the left and right of the mouse’s head). Training continued until mice ran at least 60 laps at 2 laps/min per session for three consecutive days (behavior criteria). On test day (the day after craniotomy surgery), animals were exposed to the familiar context (context A) followed by a novel context (context B). The two VR contexts (circumference = 314 cm each) had different texture backgrounds, visual objects, and reward locations. Running in the familiar context (ctx A) ended after ~60 laps or ~30 minutes, whichever came first. VR tablets were then switched off for 1-2 minutes, and a second VR context was started. After another round of ~60 laps or ~30 minutes, VR tablets were immediately switched off and urethane (750 mg/kg, subcutaneous) was administered to record urethane sleep. Subsequent 250 mg/kg booster injections were administered when movement was observed.

### Silicon probe recordings and spike-sorting

On test day, mice were head-fixed on the styrofoam ball as during training. Two 4-shank 128-channel silicon probes ^20^ coated with DiI (used for probe tracking) were positioned at specific AP and ML coordinates relative to bregma to target M2 and RSC. Probes were lowered at 1 µm/s until they reached the target depth, then allowed to settle for 45 minutes before recording started. Electrophysiological data from 256 channels was sampled at 30 KHz. After recording, LFP data was sub-sampled at 1250 Hz for further processing.

Data for M2 and RSC targeting probes were processed separately using Kilosort ^63,64^. Further manual curation was done to remove noise clusters (unusual waveforms: square components, multiple peaks, larger spread, etc.) and clusters with higher contamination (> 10 % spikes violating 2 ms refractory period). Only clusters with < 10 % contamination and a mean firing rate > 0.1 Hz were labeled good and included. Further cluster inclusion criteria were used, such as > 70% presence ratio (unit should be active in more than 70% of 10-second bins for the entire recording duration), > 40 µV amplitude, and mean firing rate > 0.1 Hz and mean firing rate < 10 Hz (to exclude interneurons) ^65^. Note that two animals were excluded from the analysis due to a small number of good excitatory units in both brain regions (total good excitatory units in each region < 30). After these inclusion criteria, we focused our analysis on data from n=2 animals.

### Spatial firing rate maps and remapping analysis

Most units were recorded in deep layers (layer 5/6) of RSC and M2. After manual curation and determining good units, we calculated the spatial firing rate map for both contexts. We first calculated the number of spikes (spike map) and total time spent (occupancy map) in 3 cm spatial bins for each trial. Epochs when the mice moved less than 2.5 cm/s and spatial bins where the animals spent < 0.5 s (across laps) were excluded. A spatial firing rate map for each lap was calculated by dividing the smoothed spike map by the smoothed occupancy map (gaussian-smoothing, σ = 1.25 bins), resulting in an L laps x P position bins matrix. We also averaged the firing rate maps across laps for each unit to calculate an average rate map, which was then used to calculate the spatial information score ^66,67^.

M2 showed a large proportion of reward or reward anticipation units ^68^ (78%, Suppl Figure 1A). Due to this, and our interest in spatially selective firing across contexts ^69^, we focused mainly on RSC units from two mice (Suppl Figure 1B,C). Pearson’s correlation between the average rate maps of individual units was used to estimate the extent of remapping between the two contexts (A-B), odd and even laps of context A (A_odd_-A_even_), and context B (B_odd_-B_even_) (Suppl Figure 1D) ^70^. The odd and even laps correlations are positive control. We observed a lower correlation between the two contexts (A-B) across all recorded units from both mice compared to the positive controls, suggesting global remapping (or more orthogonal spatial firing) across contexts ^71^.

### Sleep stage classification and Up-Down state calculation

A reference channel parked in agranular RSC Layer 5, identified using histology, and maximum power in the high-frequency (> 300 Hz) band ^72^ was used for sleep stage classification. Periods where the ratios of delta (0.5 – 4 Hz) and theta (5 – 10 Hz) power (theta/delta) remained above 5 for at least 20 seconds were classified as rapid-eye movement (REM). Periods during which theta/delta ratio remained below 2.0 for at least 20 seconds were classified as NREM (non-REM) (Suppl Figure 1E,F). Everything else was scored as wakefulness. The total duration of NREM sleep for both mice was 3633.11 s and 4398.4 s. Up-Down states were detected using spiking activity from all good units for each NREM epoch. The sum of spikes for all units was calculated in 50 ms time bins to calculate a summed PV array and a peak detection algorithm (threshold: amplitude > 7) was applied to this array to detect Up-state peaks. Edges of Up-state were defined as time points where the sum=0 (Suppl. Figure 1G). Zero population spiking times were labeled as Down states. The mean ± sem Up and Down duration was mouse1: [504.9 ± 4.5 ms, 372.9 ± 8.3 ms], and mouse2: [561.16 ± 6.3 ms, 452.24 ± 10.2 ms] (Suppl Figure 1H). Further manual checks were done to verify the results.

### PCA-based reactivation Strength and context reactivation probability

#### Reactivation strength calculation

We followed methods from Peyrache et al. ^8,28^. Individual neuron activity was first binned into 50 ms bins and z-scored to generate an N neurons x T time bins spike matrix for the entire recording duration. We then constructed an N x N pairwise correlation matrix (diagonal set to zero) from the spike matrix by computing Pearson’s correlation for both the familiar context (context A) and novel (context B) context. Each correlation matrix was factorized into orthogonal eigenvectors (principal components, PCs) with associated eigenvalue. The Marčenko-Pastur distribution was used to identify “significant components” (eigenvectors with eigenvalues exceeding the distribution’s upper bound ^73^). Interestingly for both contexts, only the first two PCs passed the threshold (total variance explained for PC1+PC2: context A: 24.61 %, context B: 23.67 %). This was true for both animals’ RSC data included in the study. The correlation matrix was then replaced by the significant PC projectors (outer product of eigenvectors) yielding a reactivation strength specific to one PC. Reactivation strength across time was calculated for urethane sleep epochs using :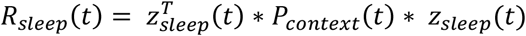, where P_context_ is a projector for a given context PC, R_sleep_(t) is reactivation strength, and z_sleep_ is the z-scored N x T spike matrix for the urethane sleep epoch. Reactivation strength using templates from contexts A and B were finally expressed as the sum of the reactivation strength of individual PCs, i.e., R_A_ = R_A_PC1_(t) + R_A_PC2_(t) and R_B_ = R_B_PC1_(t) + R_B_PC2_(t). Changing the bin width from 50 ms to 75 ms did not alter the qualitative results.

#### Context reactivation probability calculation

We compared the urethane sleep reactivation strengths calculated using familiar and novel context PCs to see if there is preferential reactivation of one context over the other in individual Up states. Each Up state detected above (duration > 300 ms) was divided into three equal parts: start, middle, and end 1/3^rd^. For each 1/3^rd^, the probability of novel context reactivation (R_B_) > familiar context reactivation (R_A_) was calculated in 4-second windows. This gave us a probability of higher reactivation strength for each context every 4 seconds. Dividing the Up-states (for example, first 100 ms, last 100 ms, and remaining segment) into different ways did not change the qualitative results reported here (Suppl Figure 4I).

## ACKNOWLEDGEMENTS

This work was supported by NIH (1R01MH125557, 1RF1NS132913), NSF (2223839).

## Contributions

R.G., O.C.G., and M.B. conceived the modeling experiments; R.G., O.C.G., and J.E.D. conducted the simulations; R.G. analyzed the simulated data; S.K. and B.L.M. conceived the rodent experiments; S.K. conducted the rodent experiments and collected the empirical data; R.S. processed and analyzed the empirical data; R.G. and M.B. wrote the first draft of the manuscript; all authors reviewed and edited the manuscript.

## ETHICS DECLARATIONS

### Animal Welfare

All experimental protocols were approved by the Institutional Animal Care and Use Committee of the University of California, Irvine.

### Competing interests

The authors declare no competing interests.

## Supplementary Data Figures

**Supplementary Figure 1.**
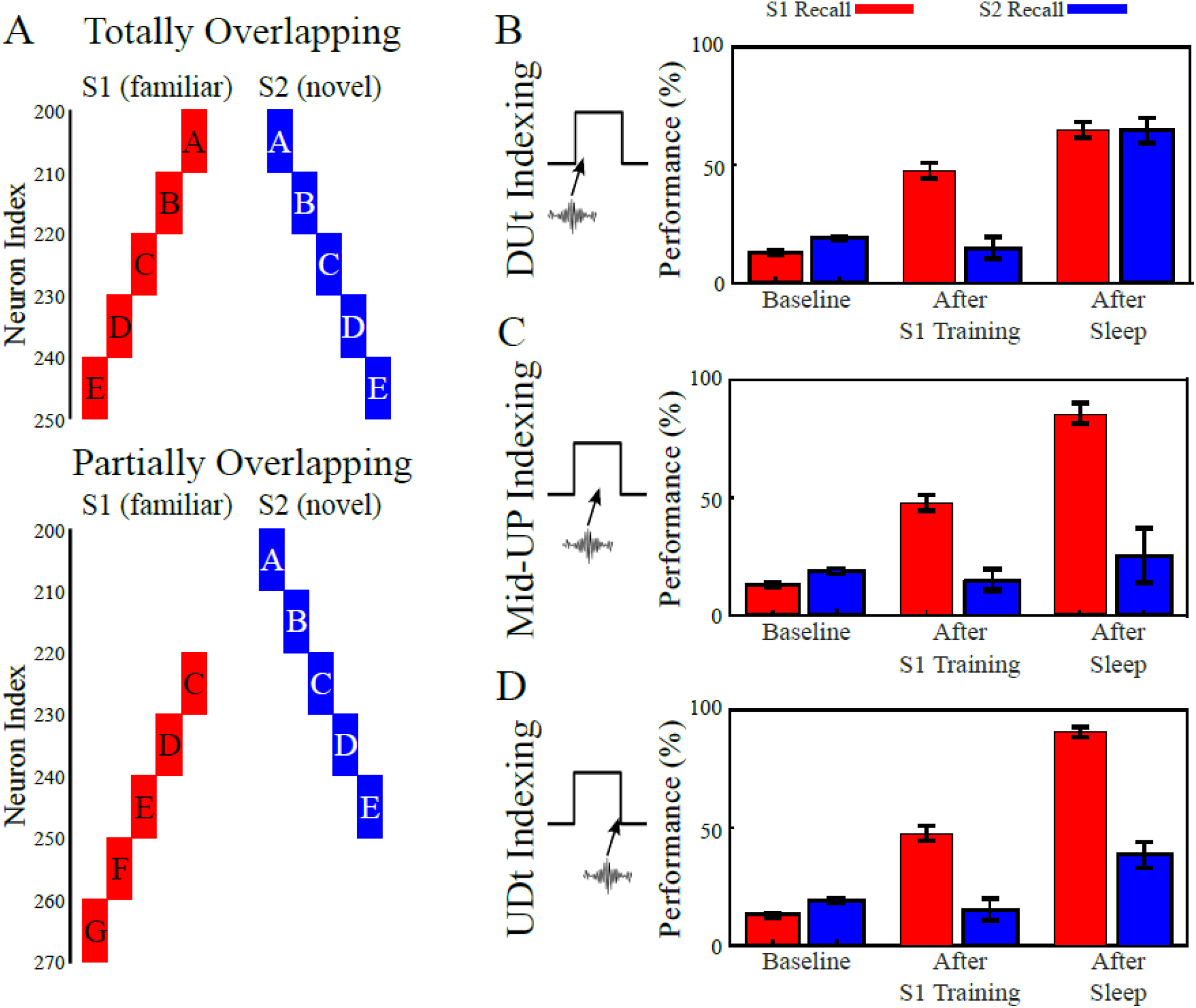
Partial Overlap Simulations. **(A)**. Schematic of the familiar (S1; red) and novel (S2; blue) memory sequence representations used for the model in the case of the original totally overlapping case used throughout the paper (top) and of the partially overlapping case used here (bottom). **(B-D)** Bar plots showing familiar S1 (red) and novel S2 (blue) recall performance at baseline, after S1 training, and after sleep for the partially overlapping memories with DUt Indexing (B), Mid-UP Indexing (C), and UDt Indexing (D). Error bars denote standard deviation.

**Supplementary Figure 2.**
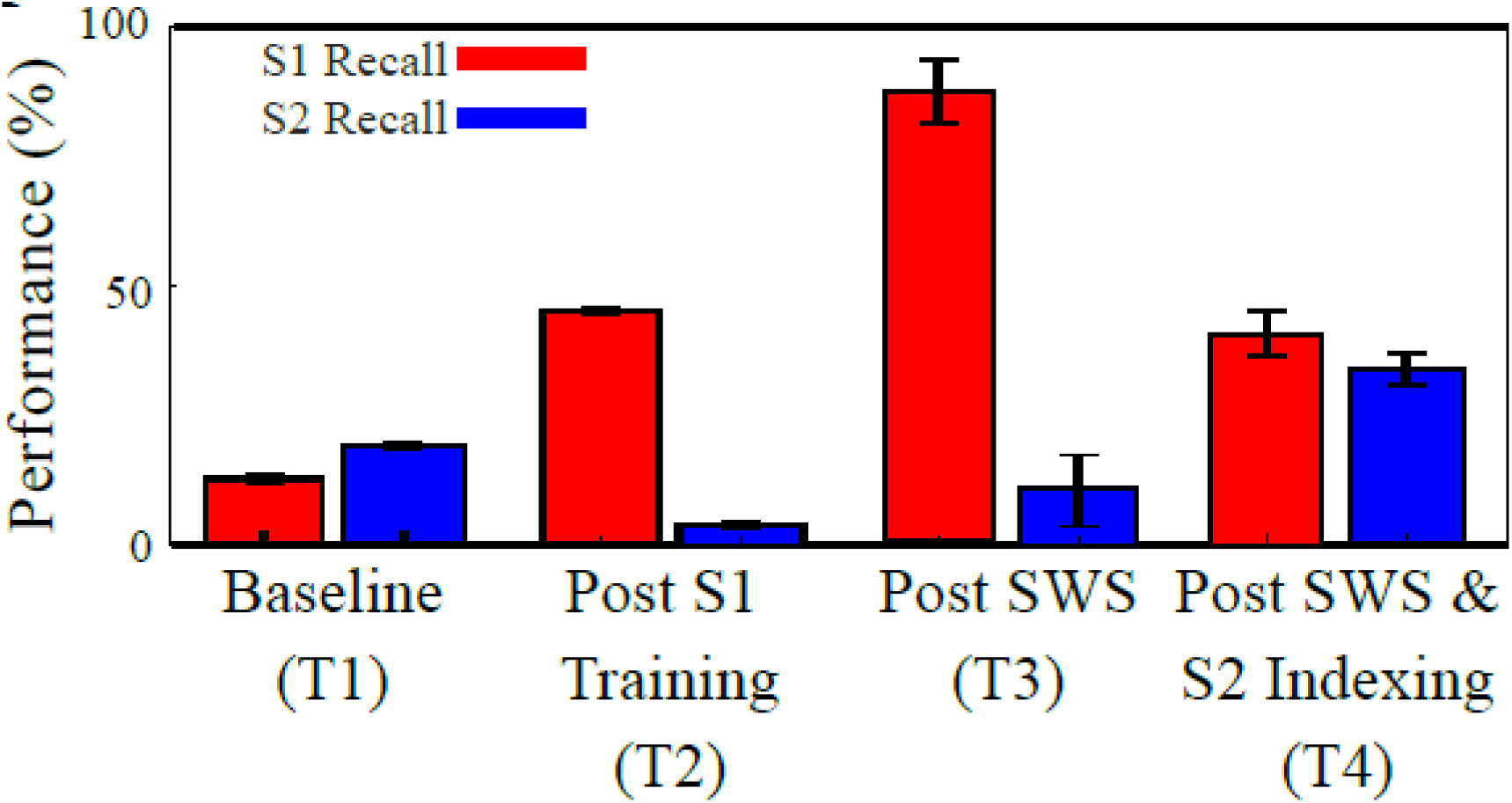
Bar plots showing S1 (red) and S2 (blue) recall performance at baseline (T1), after 250s of S1 training (T2), after 2000s of SWS (T3), and after another 3500s of SWS with S2 indexing occurring on the DUt of each Up state. Error bars denote standard deviation.

**Supplementary Figure 3.**
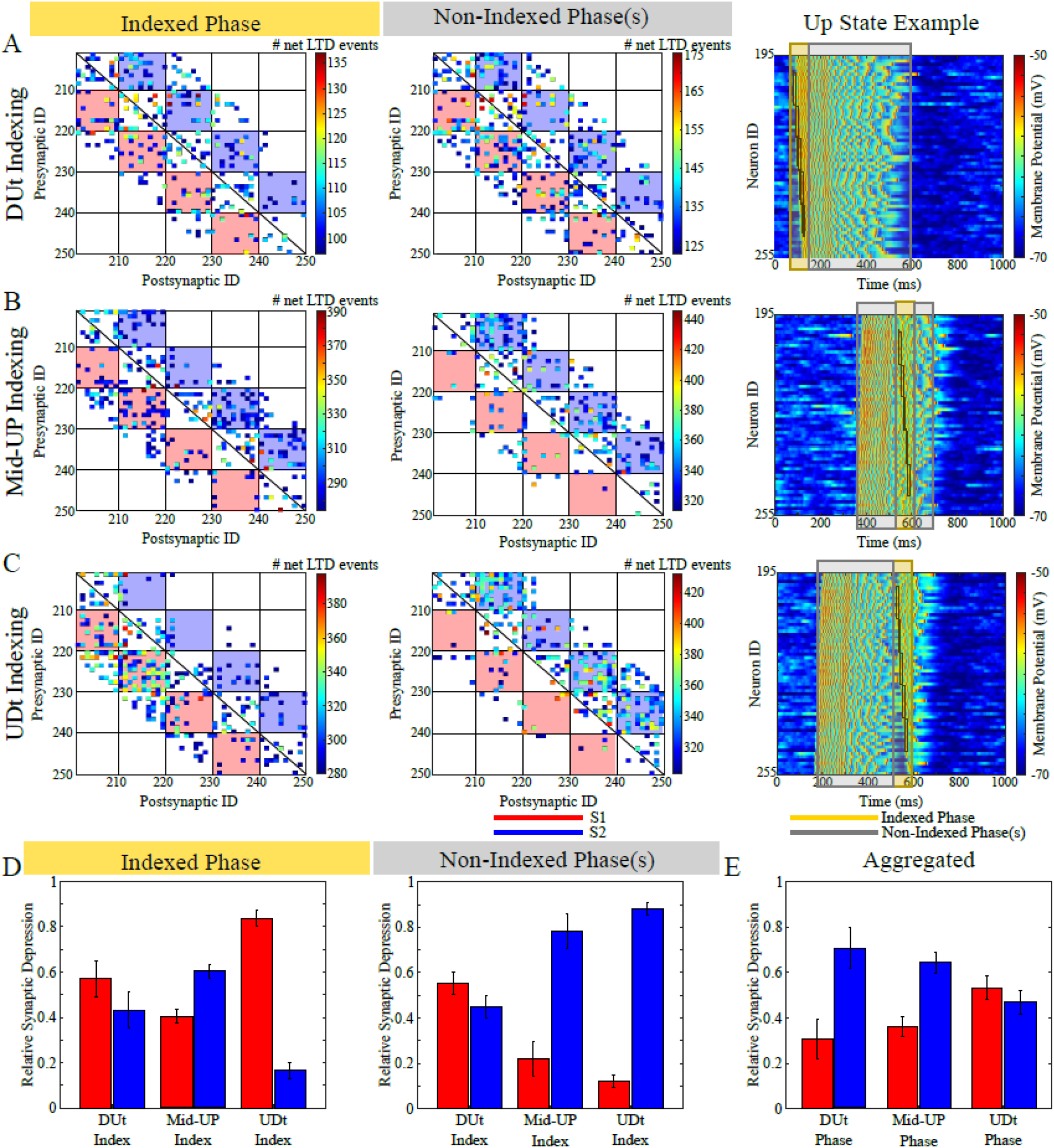
Net synaptic depression. **(A-C)** Heatmaps display the number of UP states (approximately 450 total UP states per simulation) in which a synapse underwent a net LTD event. The heatmaps illustrate events occurring during the phase of the UP state when the artificial index was applied (Indexed Phase; left) or during the phase(s) of the UP state without indexing (Non-Indexed Phase(s); middle). In three independent experiments, the index was applied at either (A) the DUt, (B) the Mid-UP, or (C) the UDt. The background red/blue colors indicate synapses supporting the transitions for S1 and S2, respectively. The Up state example (right) shows a single Up state for each of the three indexing conditions, depicting the indexed phase (yellow shading), the non-indexed phases (gray shading), and the applied index (black boxes). The colormap represents the number of net LTD events for each synapse. To avoid visual clutter, only the top 30% of synapses are shown. **(D)** Bar plots show the relative amount of synaptic depression for S1 (red) versus S2 (blue) during the Indexed Phase (left) and Non-Indexed Phase(s) (right). Relative synaptic depression was calculated by summing the number of net LTD events in the red/blue squares for S1/S2, respectively, and normalizing these values by the total number of events for both memories (i.e., the sum of events in all colored squares). This normalization accounts for differences in population-level firing rates across different Up state phases. **(E)** Bar plots display relative synaptic depression across the three phases (DUt, Mid-UP, and UDt), with data aggregated across all three simulation conditions. Error bars in (D-E) represent standard deviations.

**Supplementary Figure 4.**
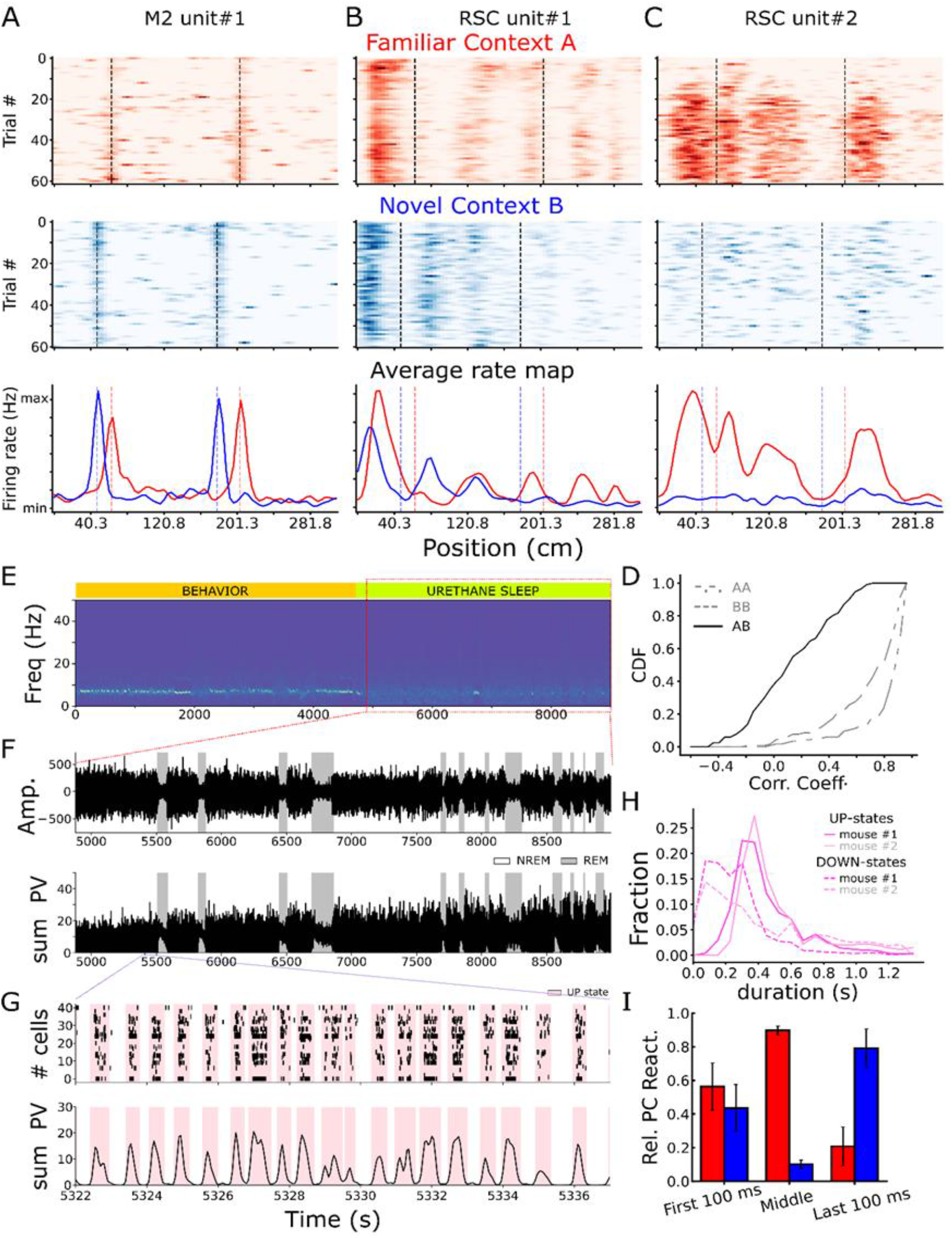
Spatial firing ratemaps for familiar (first row, red) and novel context (second row, blue), and average firing rate maps (third row) for single-units in M2 **(A)**, and RSC **(B**,**C). (D)** Cumulative distribution plot for Pearson’s correlation coefficient between contexts (AB, solid black line), odd and even laps of each context (A_odd_-A_even_ (dotted-dashed gray line), B_odd_-B_even_ (gray dashed line). **(E)** spectrogram for the entire behavior duration. Notice high power in the theta range (6 – 10 Hz) during behavior epochs (orange), and a lack of high theta power in urethane epochs (green). **(F)** Delta oscillation (0.1 – 4 Hz) signal (µV, top) and sum population vector (PV, 50 ms bins, bottom) for ~3600 s long urethane sleep epochs. Gray highlighted zone mark REM epochs. **(G)** Zoom-in on 15 s NREM epoch showing detected UP (pink) and DOWN states in spike raster (top) and sum population vector (PV, bottom). **(H)** Distribution of UP (solid line) and DOWN (dashed line) state duration for both animals. **(I)** Mean ± sem familiar (red) and novel (blue) context relative PC reactivation for first 100 ms, last 100 ms, and remaining segment of UP-states for mouse #1.

**Supplementary Figure 5.**
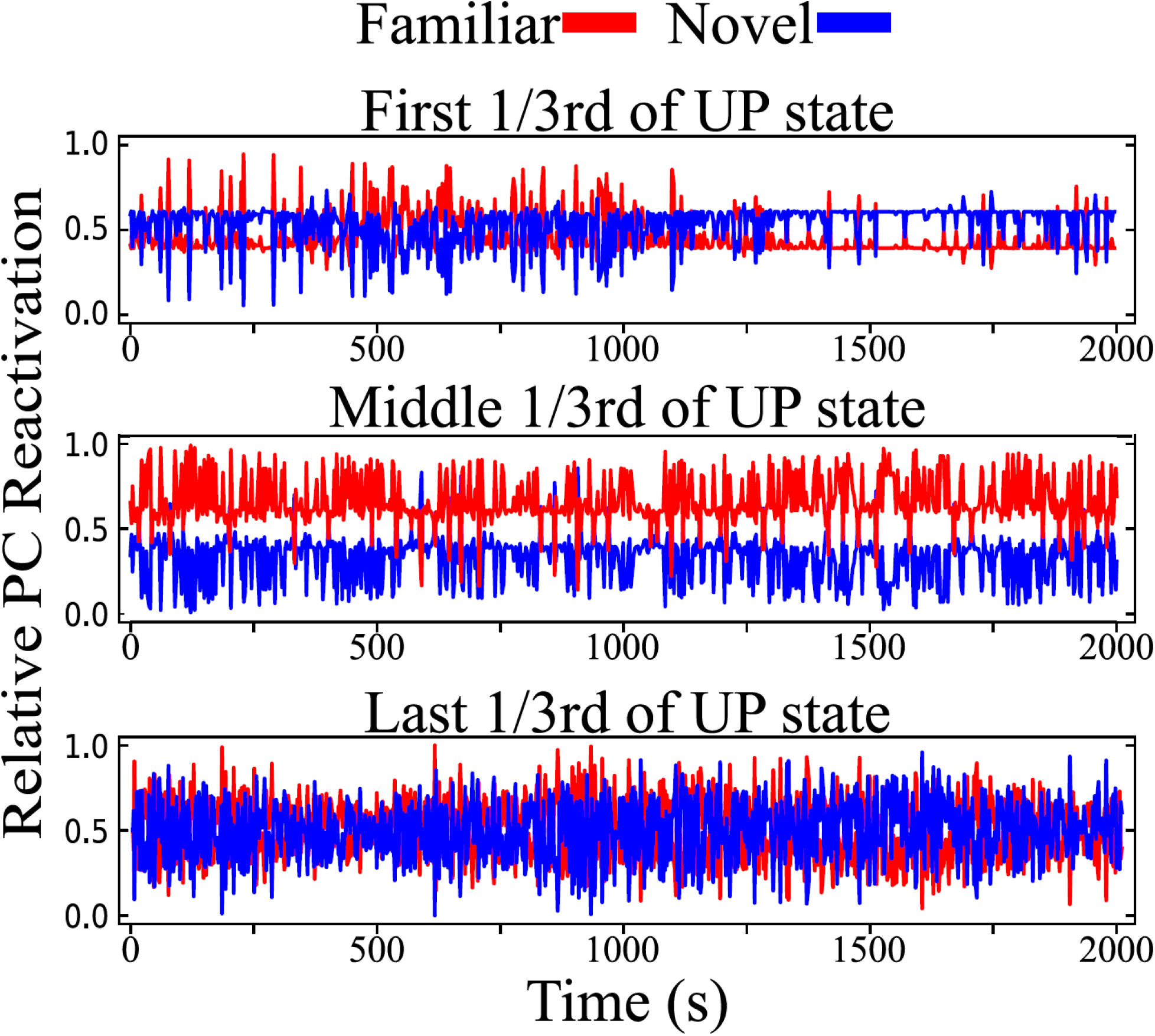
Relative PC reactivation in model during SWS. The PCA-based reactivation measure applied to rodent data (Figure 4A) was applied to the simulation with stochastic indexing (Figure 4D). Partially overlapping memory representations were used as this measure could not discriminate between fully overlapping representations due to high similarity between correlational structures.

## Notes

### Competing Interest Statement

The authors have declared no competing interest.

### Summary of Updates

Expanded introduction and discussion to more explicitly include the inspiration from the early AI / Connectionist literature of the mid-1900s as well the the relevance to modern machine learning.

